# *Bacillus thuringiensis* targets the host intestinal epithelial junctions for successful infection of *Caenorhabditis elegans*

**DOI:** 10.1101/338236

**Authors:** Liting Wan, Jian Lin, Hongwen Du, Alejandra Bravo, Mario Soberón, Donghai Peng, Ming Sun

## Abstract

Pathogenic bacteria use different strategies to infect their hosts including the simultaneous production of pore forming toxins and several virulence factors that help to synergize their pathogenic effects. However, how the pathogenic bacteria are able to complete their life cycle and break out the host intestinal barrier is poorly understood. The infectious cycle of *Bacillus thuringiensis* (Bt) bacterium in *Caenorhabditis elegans* is a powerful model system to study the early stages of the infection process. Bt produces Cry pore-forming toxins during the sporulation phase that are key virulence factors involved in Bt pathogenesis. Here we show that during the early stages of infection, the Cry toxins disrupt the midgut epithelial tissue allowing the germination of spores. The vegetative Bt cells then trigger a quorum sensing response that is activated by PlcR regulator resulting in production of different virulence factors, such as the metalloproteinases ColB and Bmp1, that besides Cry toxins are necessary to disrupt the nematode epithelial junctions causing efficient bacterial host infection and dead of the nematode. Overall our work describes a novel mechanism for Bt infection, targeting the epithelial junctions of its host midgut cells.

**Author summary:** The entomopathogenic bacteria *Bacillus thuringiensis* (Bt) are used worldwide as biopesticides due to their insecticidal properties. Crystal proteins (Cry) produced by Bt during the sporulation phase of growth are mainly responsible for their insecticidal properties. The infection process of Bt includes three successive steps, virulence, necrotrophic, and sporulation processes. During the virulence process, after ingestion by the susceptible hosts, the Cry toxins form pores in the apical membrane of intestinal cells, inducing favorable conditions for bacterial spore germination. Vegetative bacteria multiply in the host and coordinate their behavior by using the quorum sensor regulator PlcR, which leads to the production of virulence factors allowing the bacteria to kill the host. However, how the bacteria are able to disrupt the host intestinal barrier during the early stages of infection remains unknown. Here we show that Bt employs the nematicidal Cry toxins and additional virulence factors controlled by the PlcR regulon to disrupt the intestinal epithelial junctions of *C. elegans* at the early stages of infection allowing that Bt bacteria complete its life cycle in the worms. Our work provides new insights into the pathogenesis of Bt, and highlights the importance of breaking down host epithelial junctions for a successful infection, a similar mechanism could be used by other pathogens-host interactions since epithelial junctions are conserved structures from insects to mammals.

## INTRODUCTION

Animals encounter a myriad of chemical or biological dangers, including pathogens, in the environment. The epithelium tissue naturally serves as a physical barrier to protect themselves from detrimental agents [1]. However, pathogenic bacteria evolved to avoid the defense mechanisms and to invade their host [2,3] completing their infection lifecycle [4,5]. Indeed, it has been reported that many pathogens disrupt the host intestinal barrier functions during infection, such as the human opportunistic pathogen *Escherichia coli* [6], an etiological agent of human adult periodontitis *Porphyromonas gingivalis* [7], and *Salmonella* enterica serovar *typhimurium* [8], among others. However, there have been very few studies addressing the detailed mechanism of how pathogenic bacteria break through the intestinal barrier of their hosts during the early stages of infection.

The Gram-positive bacteria *Bacillus thuringiensis* (Bt) are ubiquitous spore-forming bacteria that belong to the *Bacillus cereus* (Bc) group, which also encompasses *B. cereus* (a human opportunistic pathogen), and *B. anthracis* (the etiological agent of anthrax in mammals) [9]. During sporulation, Bt produces parasporal crystal proteins (Cry) that have specific toxicity to different insects, nematode, mites, and protozoa species. [10,11]. The Cry proteins are recognized as pore-forming toxins that disrupt the epithelial cells by osmotic shock [12]. In addition, Bt produces several other virulence factors important for its pathogenicity in insects [13]. For example, the enhancin protein was reported to facilitate the disruption of the peritrophic membrane, while the InhA metalloprotease helps Bt and Bc to escape from innate immune system. In the case of Bc it was shown that InhA metalloprotease induces alterations in the macrophage membrane [14-16].

As a pathogen, Bt could complete a full infection life cycle in both insects and nematodes hosts [13,17]. Overall, the infection process of Bt includes three successive steps, that involve virulence, necrotrophic, and sporulation processes [18]. During the virulence process, after ingestion by susceptible hosts, Cry toxins form pores in the host intestinal cell membranes, inducing favorable conditions for spore germination. Vegetative bacteria multiply in the host and coordinate their behavior by using the quorum sensor PlcR regulator, which leads to the production of virulence factors allowing the bacteria to kill its host [19,20]. After the host death, the quorum sensor NprR regulator controls the expression of a set of genes encoding degradative enzymes, allowing the vegetative bacteria cells to use the insect cadaver and to survive necrotrophically [21]. Finally, the Rap-Phr system activates a phosphorylation cascade leading to the induction of the sporulation regulator Spo0A and to the commitment of part of the bacterial population to produce Cry protein and sporulate [18,19]. Although some steps related to Bt infection have been studied, it is still not known how the bacteria disrupt the intestinal barrier of the host during the early stages of infection.

The infectious cycle of Bt in the nematode *C. elegans* is a powerful model system to study the pathogenesis under natural conditions [17,22]. Indeed, several virulence factors of Bt besides Cry toxins were shown to contribute to the nematodes infection, including the metalloproteases Bmp1, ColB, and chitinase [22-24]. Especially, the Bmp1 and ColB exhibit intestinal tissue degradation activity to enhance the toxicity of Bt. In this study, using Bt*-C. elegans* as a model, we describe the detailed mechanism of how Bt breaks out the intestinal barrier of the host. We report that the intestinal epithelial junctions of *C. elegans* are targeted by Bt to invade the worms at the early stages. Also, we show the genetic basis and regulation of the virulence factors involved in this process. Our findings point out a crucial role of PlcR regulator and the virulence factors, Bmp1 and ColB in disrupting the nematode epithelial junctions. Overall, we provide new insights into the pathogenesis of Bt, and highlight the importance of breaking down host epithelial junctions for successful infection.

## Results

### Nematicidal Bt bacteria completed their life cycle in *C. elegans* destroying host internal structures

We previously described the complete lifecycle of the nematicidal Bt strain YBT-1518 in *C. elegans* [22]. To confirm whether this was a special case of that particular Bt strain or an extensive phenomenon, we tested the pathogenicity in *C. elegans* of other five wild-type Bt strains [25] and one recombinant Bt strain (BMB0215) [26]. All of these Bt strains are nematicidal expressing the major classes of nematicidal Cry toxins such as Cry5, Cry6, Cry14, Cry21 and Cry55 (Table S1) [27]. As negative controls we used a non-nematicidal Bt strain YBT-1520 [28] that has high toxicity against lepidopteran insects such as *Plutella xylostella* and *Helicoverpa armigera*, and an acrystalliferous mutant of Bt strain BMB171 that encodes no *cry* genes and is nontoxic to the nematodes or to the lepidopteran insects [29]. The microscopy observations showed that the spores of all Bt strains escape from being destroyed in the pharyngeal grinder and reached the intestine of the nematodes (data not shown). The spores of the seven Bt strains that have nematicidal activity, were able to germinate and developed into vegetative cells after 24 h post infection (hpi). After 72 hpi the body of these infected nematodes were filled with Bt vegetative cells; while the tissues and organs of infected nematodes were degraded, showing Bt vegetative cells wrapped inside the nematode epidermis (Fig. 1A), which is a phenotype known as “bag of bacteria” (“Bob”) [30]. In contrast, the nematodes infected with non-active Bt strains BMB171 and YBT1520 showed spores in the gut but no vegetative Bt cells were observed in intestinal tissue of the nematodes and neither showed “Bob” phenotype (Fig. 1A).

**Fig. 1.**
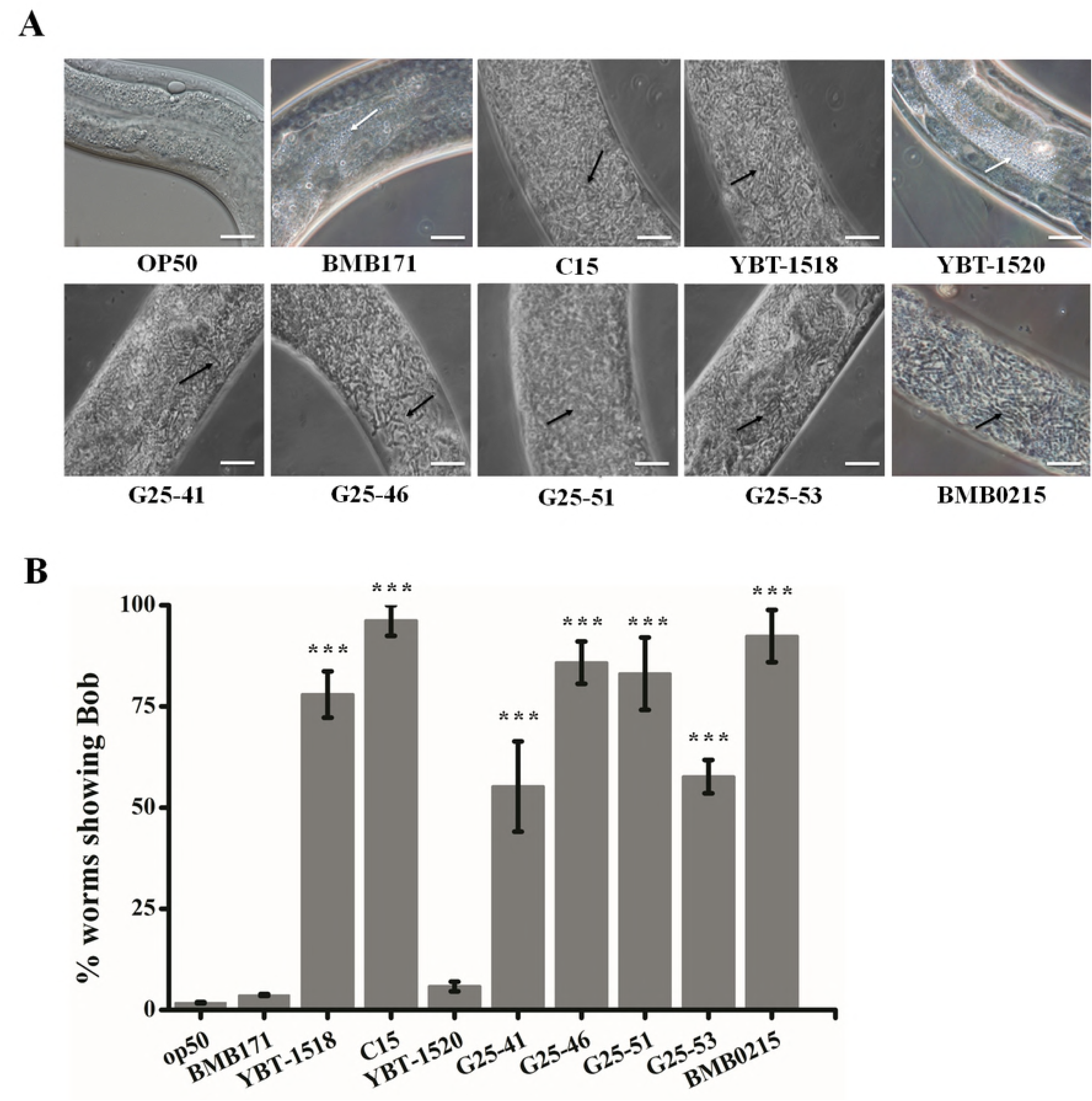
Nematicidal Bt bacterium destroys the internal structures of the nematodes. (A) The pathogenicity behavior of selected Bt strains in N2 worms was assessed by the “Bob” phenotype assay. Representative images of worms treated with different Bt strains after 72 hpi are shown. Scan bars represent 10 μm. (B) Analysis of the percentage of worms that showed “Bob” phenotype after treatment with each Bt strain. The mean and standard error deviations of three independent experiments are shown. Triple asterisks indicate a significant difference (*t*-test, *p*<0.001) and no labeled indicates no significant difference between these treatments compared with that of *E. coli* OP50 strain. BMB171 is an acrystalliferous Bt strain. YBT-1520 is a Bt strain active against Lepidopteran insects, non-active against nematodes.

The percentage of worms that showed “Bob” phenotype after treatment with each Bt strain was quantified indicating that most of the worms infected with the nematicidal strains formed “Bob” phenotype, ranging from 55.19 ± 11.16 % (infected by Bt strain G25-41) to 96.20 ± 3.78% (infected by Bt strain C15) (Fig. 1B). All worms in the two control groups were healthy (Fig. 1B). Thus, we concluded that the nematicidal Bt strains could complete their full life cycle in *C. elegans*, and shared common pathogenic features destroying the worm’s internal structures.

### Epithelial junction related genes of *C. elegans* are upregulated during Bt infection

To explore the detailed mechanism of how nematicidal Bt destroys the intestine structures of nematodes, we previously analyzed the *C. elegans* proteome after infection with a nematicidal Bt strain in comparison to a non-nematicidal Bt strain [31]. Interestingly, these studies showed that three intestinal epithelial junction components related proteins AJM-1, DLG-1, and LET-413 were significantly up-regulated after nematicidal Bt infection [31]. The DLG-1 protein could physically interact with AJM1 in the *C. elegans* epithelial junction, while the DLG-1/AJM-1 complex is targeted to apical junctions by LET-413, which may be linked to the cell membrane by claudin-like transmembrane proteins such as VAB-9 or CLC-1 [32]. These data indicate that the epithelial junction related genes may be involved in the response to Bt infection.

To confirm this hypothesis, we analyzed the expression of the epithelial junction related genes *clc-1, ajm-1, dlg-1*, and *let-413* by qPCR in *C. elegans* worms exposed to the six different nematicidal Bt strains in comparison with two non-nematicidal Bt strains. Also, worms fed with *E. coli* OP50 strain were used as negative control. The results showed that the epithelial junction related genes *clc-1, ajm-1, dlg-1*, and *let-413* were significantly up-regulated (*t-*test, *p*< 0.01) in animals infected with all the nematicidal Bt strains in contrast with the worms feed with *E. coli* OP50 strain or with the non nematicidal Bt strains (Fig. 2). It is important to mention that no significant differences (*t-*test, *p*> 0.05) were found in worms fed with the two non-nematicidal Bt strains in comparison to the treatment with the negative control *E. coli* OP50 strain (Fig. 2). This information is consistent with the previously proteomic analysis [31] supporting that the epithelial junction related genes of *C. elegans* were significantly upregulated during nematicidal Bt infection.

**Fig. 2.**
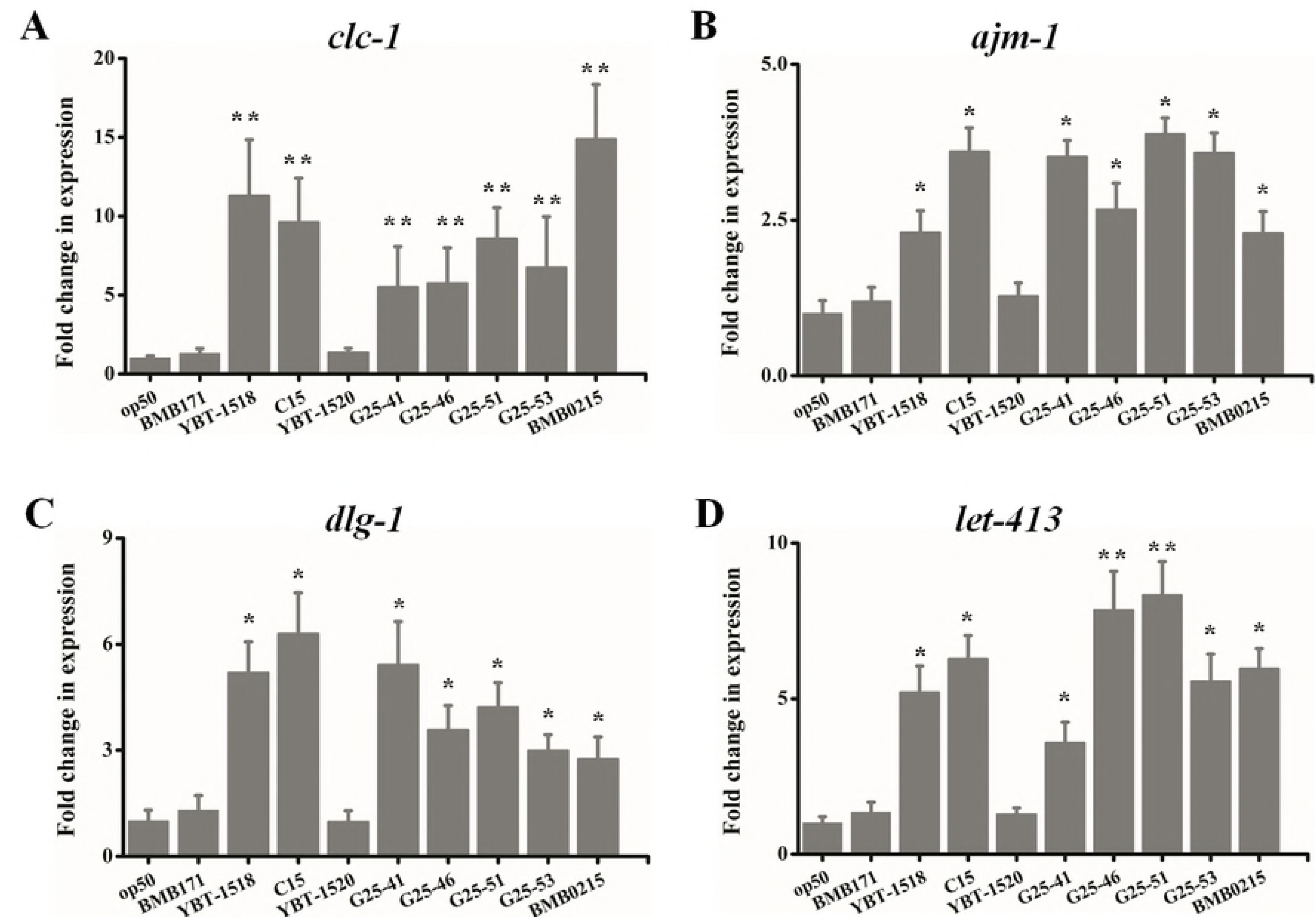
Several epithelial junction related genes of *C. elegans* are significantly upregulated after Bt infection. The expression levels of *C. elegans* epithelial junction related genes were monitored by qPCR after exposure to different Bt strains for 10 h. Asterisks indicate significant difference (Student’s *t*-test, *, *p*<0.05; **, *p*<0.01) and no label indicates any significant difference between these treatments compared with that of *E. coli* OP50 strain. BMB171 is an acrystalliferous Bt strain. YBT-1520 is a Bt strain active against Lepidopteran insects, non-active against nematodes.

### Nematicidal Bt strains disrupt the intestinal epithelial junctions of *C. elegans* during infection

The intestine of *C. elegans* is comprised of 20 large epithelial cells that are mostly positioned as bilaterally symmetric pairs to form a long tube around the midgut lumen. The epithelial junctions serve as link of all the intestinal cells [32]. The DLG-1/AJM-1 complex is considered as one of the three major molecular complexes at the apical surface of epithelial cells in *C. elegans* [33]. Host epithelial junctions could serve as targets for the invasions of microorganisms [5,34]. Therefore, we hypothesized that the epithelial junction may be a target for Bt to destroy the intestine structures of the worms.

To confirm this hypothesis, we used the GFP labeled transgenic worms FT63(DLG::GFP) as a DLG-1/AJM-1 complex marker to visualize the integrity of *C. elegans* epithelial junctions during infection with Bt strains. The FT63(DLG::GFP) worms were fed with spores and toxin mixtures of the seven nematicidal Bt strains and observed after 24 hpi, a time point where all worms are still alive. Animals treated with non-nematicidal Bt strains or *E. coli* OP50 strain at the same conditions were used as negative controls. All treated worms were monitored and photographed under the fluorescent microscope. Normally, the GFP fluorescence of healthy FT63(DLG::GFP) worms is observed intact, just like a ladder. In contrast, when the epithelial junction is disrupted, the GFP fluorescence becomes blurry or fragmented. The observations of Fig. 3 showed that all nematicidal Bt strains caused severely disruption of *C. elegans* epithelial junctions. In contrast, the non-nematicidal Bt strains BMB171 and YBT-1520, showed that the fluorescence of the FT63(DLG::GFP) worms was similar to the animals that were fed with *E. coli* OP50 (Fig. 3). To quantify the damage in the epithelial junction’s, we observed 100-120 worms randomly selected for each group, and the proportion of worms with disrupted epithelial junctions was calculated. The results show that most of the worms infected by the nematicidal Bt strains have disrupted epithelial junctions, ranging from 50.52 ± 3.57 % (infected with Bt strain G25-41) to 88.95 ± 7.23% (infected with Bt strain BMB0215). In contrast, the non-nematicidal Bt strains did not significantly (*t-test, p*> 0.05) disrupt the epithelial junction of FT63(DLG::GFP) worms when compared with the *E. coli* OP50 control strain (Fig. 4). These results show that the nematicidal Bt disrupt the intestinal epithelial junctions of *C. elegans* during infection.

**Fig. 3.**
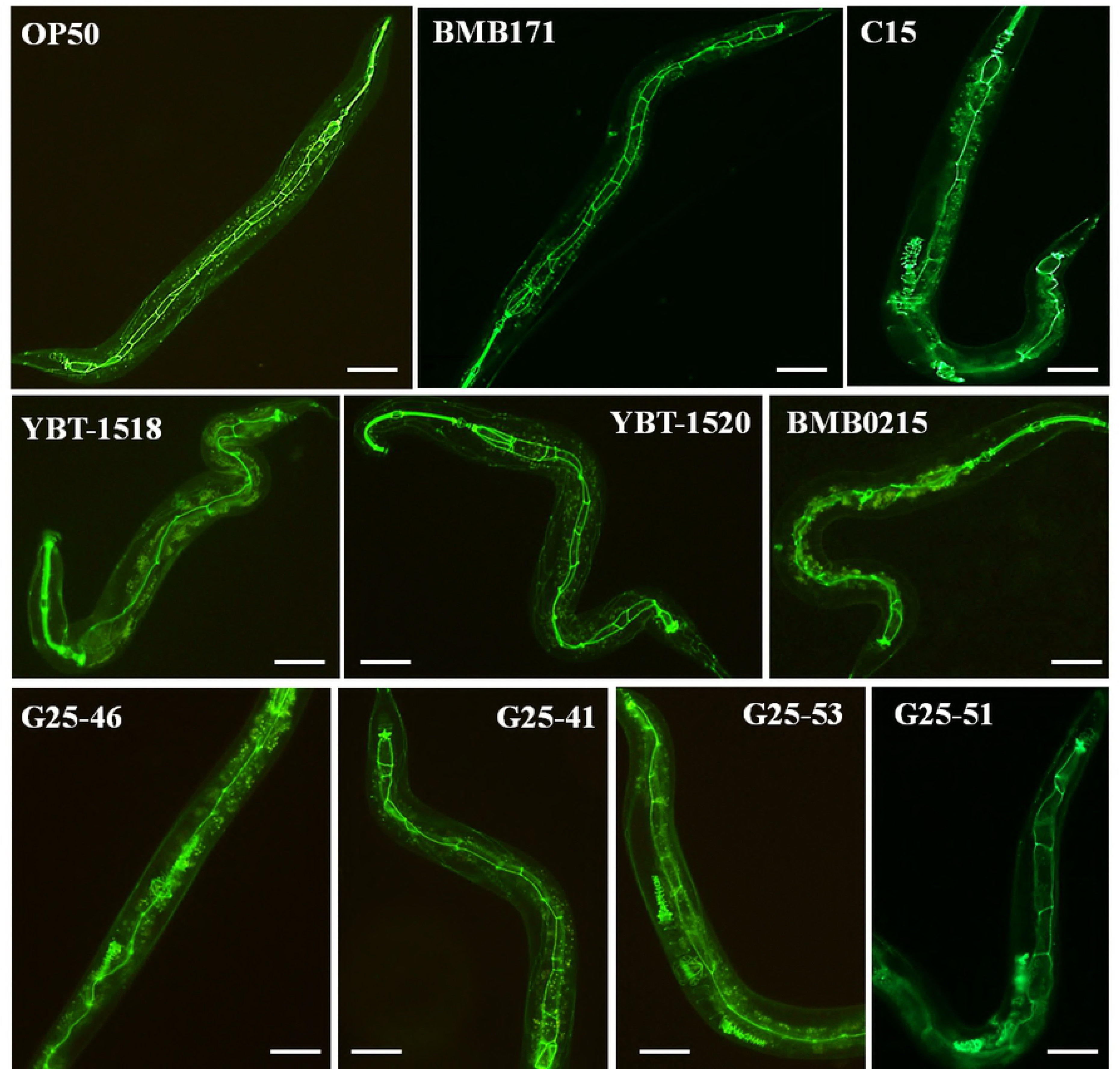
The nematicidal Bt strains disrupts the intestinal epithelial junctions of *C. elegans.* The transgenic worms FT63(DLG::GFP) were fed with selected Bt strains. The intestinal epithelial junctions of worms were observed under fluorescent microscope at 100X magnification after 18 hpi. Representative images of worms that were treated by each Bt strain are shown. The scan bars represent 20 μm.

**Fig. 4.**
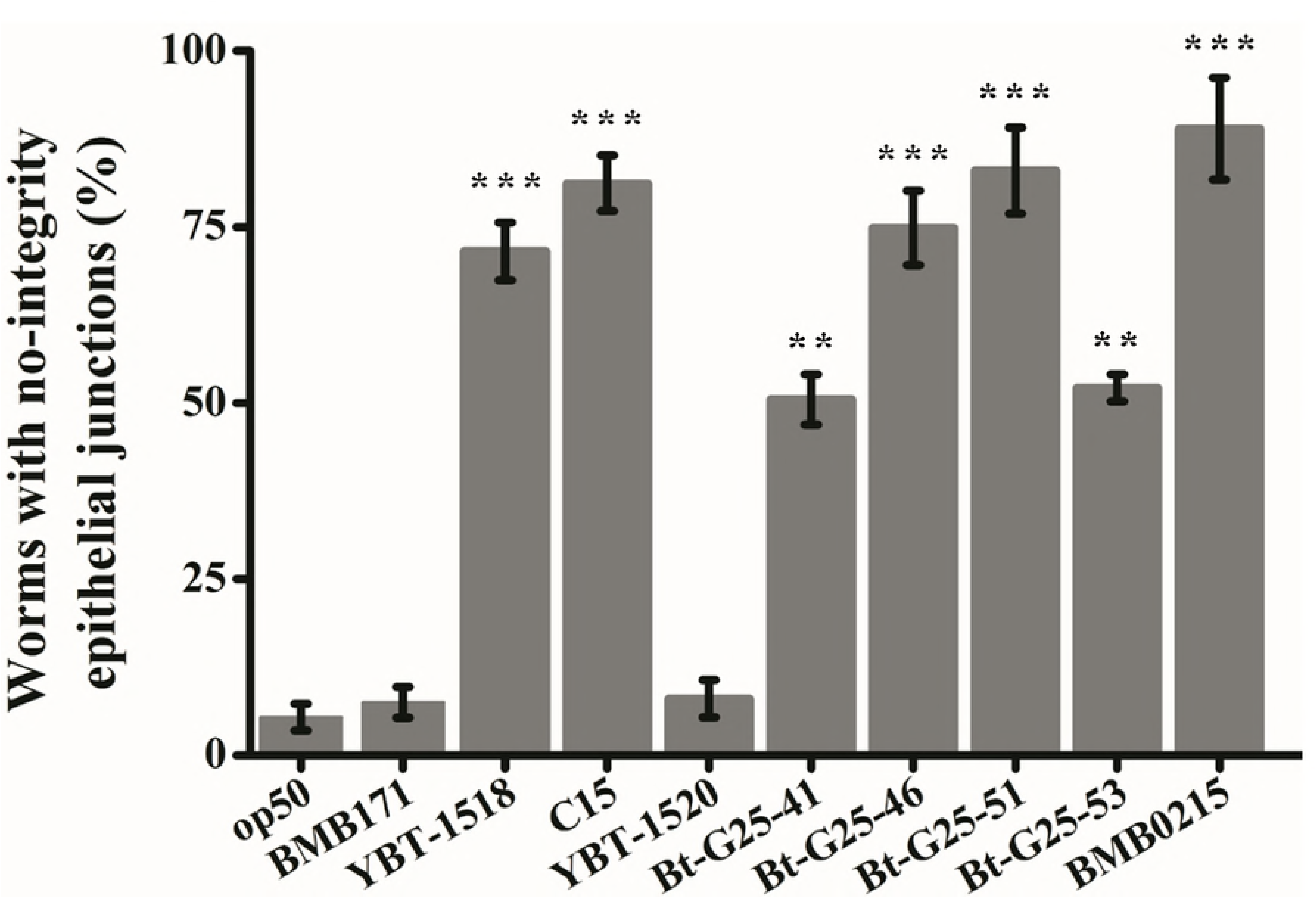
Extent of disruption of epithelial junctions caused by different Bt strains. The data are expressed as the percentages of worms that showed disrupted epithelial junctions after 18 hpi with each Bt strain. The mean and standard error of three independent experiments are shown. Asterisks indicate significant difference (Student’s *t*-test, **, *p*<0.01; ***, *p*<0.01) and no label indicates no significant differences between these treatments compared with that of *E. coli* OP50 strain.

### *C. elegans* susceptibility to Bt is associated with epithelial junction integrity

Cell-cell junctions play a critical role in cell polarity and organogenesis. In addition to these fundamental functions, they also contribute to the formation and establishment of a physical barrier against pathogens invasion [35]. Therefore, disrupting the intestinal epithelial junctions of *C. elegans* seems to be one of the Bt strategies to infect the worms. To test this hypothesis, we analyzed the nematicidal activity of all selected nematicidal Bt strains against the wild type worms N2 and against the epithelial junction’s deficient mutant worms *clc-1(ok2500)*, in which the epithelial junction related gene *clc-1* was knockout. Considering that the selected nematicidal Bt strains show high toxicity to the nematodes, we adjusted the medium lethal concentration (LC_50_) of each strain in order to have 50% mortality of the wild type worms N2. The non-nematicidal Bt strain BMB171 was used as a negative control, using a dose of 10^6^ spores/µl which is equally to the highest dose for selected nematicidal Bt strains in this assay. The results of the mortality assay showed that the mutant worms *clc-1(ok2500)* were significantly more sensitive to all the nematicidal Bt strains compared with that of the wild type N2 worms (*t*-test, *p* < 0.05) (Fig. 5A).

**Fig. 5.**
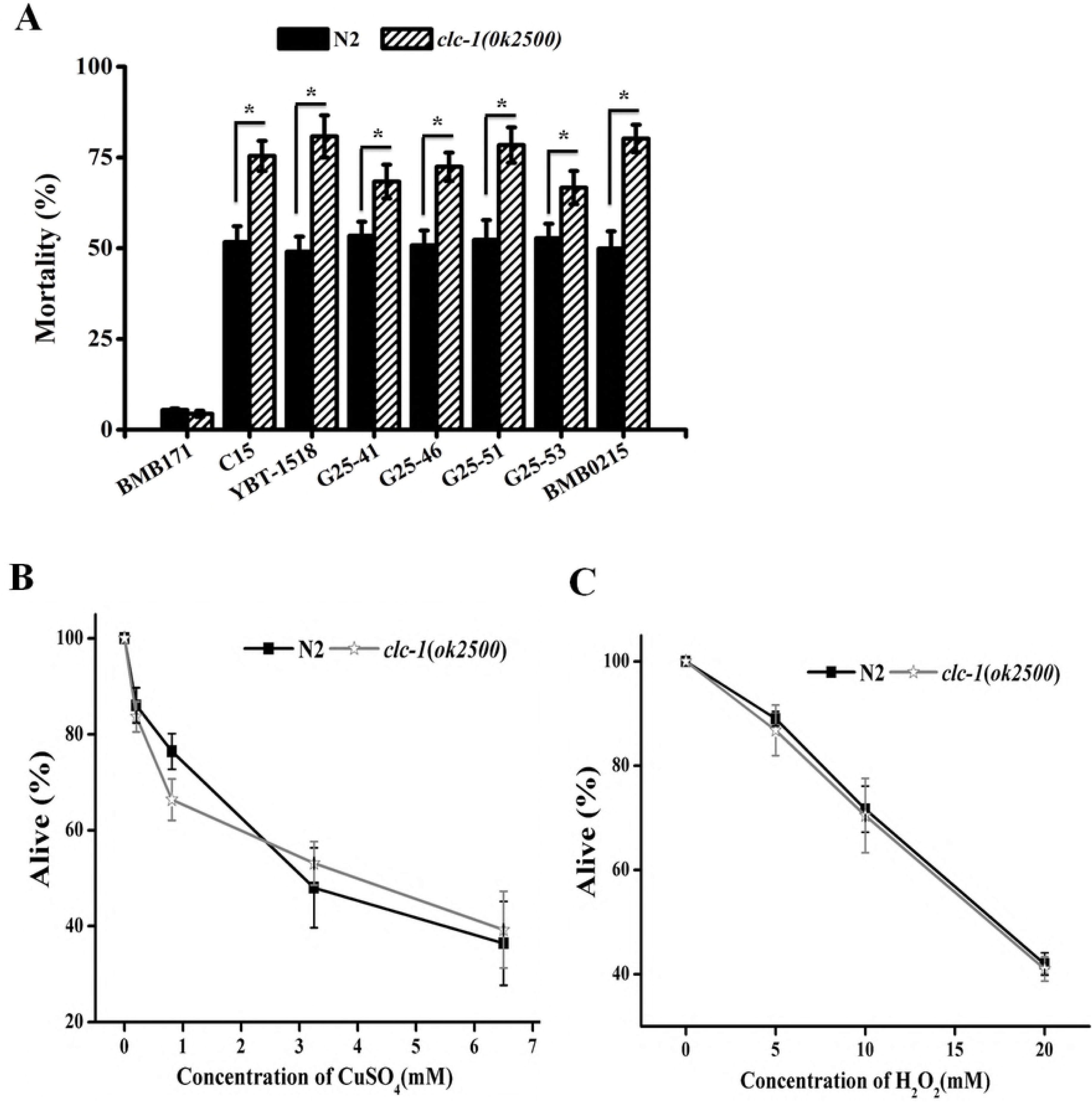
Toxicity of Bt to *C. elegans* is associated with the low epithelial junction integrity. (A) The mortality assay of nematicidal Bt strains against the wild type worms N2 or the epithelial junction mutant worms *clc-1(ok2500)*. The mean and standard error of three independent experiments are shown. (B and C) The mortality assay of the copper sulfate and hydrogen peroxide against the wild type worms N2 or the epithelial junction mutant worms *clc-1(ok2500)*. The mean and standard error of three independent experiments are shown. Asterisks indicate significance as determined by a Student’s *t*-test (*p*<0.05). No labeled indicates no significant difference.

We also measured the sensitivity of wild type and mutant nematodes to copper sulfate (Fig. 5B) and hydrogen peroxide (Fig. 5C) as controls of non-specific toxicity. There were no significant differences to these treatments between mutant worms *clc-1(ok2500)* compared with that of the wild type N2 worms (*t*-test, *p*>0.05). Our results show that the intestinal epithelial of *C. elegans* is a physical barrier against the infection of Bt.

### The Bt spore development and nematicidal Cry toxin are required for the disruption of the epithelial junction in the nematode

Cry toxins have been shown to be the main Bt virulent factor to kill the nematodes [26]. Therefore, we determined the importance of the nematicidal Cry toxins in the disruption of the epithelial junctions after Bt infection. We fed FT63(DLG::GFP) worms with 10 μg/μl of pure nematicidal Cry5Ba, Cry6A, and Cry21Aa toxins, respectively. The results showed that after 24 hpi a small proportion of the worms showed disruption of their epithelial junctions (Fig. 6), ranging from 10.0 ± 1.0 % when treated with Cry6Aa toxin to 33.0 ± 5.0% when treated with Cry5Ba toxin, compared with the disruption observed in the presence of purified spores of BMB171, that do not produce Cry toxins, together with Cry toxins at the same concentration as above (10 μg/μl), the epithelial junctions of most of the worms were destroyed, ranging from 51.7 ± 5.9% for BMB171 spores plus Cry6Aa to 91.6 ± 8.3% for BMB171 spores plus Cry5Ba. The control included worms that were fed with purified spores of BMB171, that showed very few worms with disrupted epithelial junctions (Fig. 6). As an additional control we used inactivated BMB171 spores to fed the worms plus different Cry toxins. Under these conditions there were few worms with disrupted epithelial junctions compared with those worms fed with live spores plus different Cry toxins, and the amount of affected worms were similar to those treated with pure Cry toxins (Fig. 6). These data indicate that live spores, not the components of spores, are important for the disruption of the nematode epithelial junctions induced by the Bt strains.

**Fig. 6.**
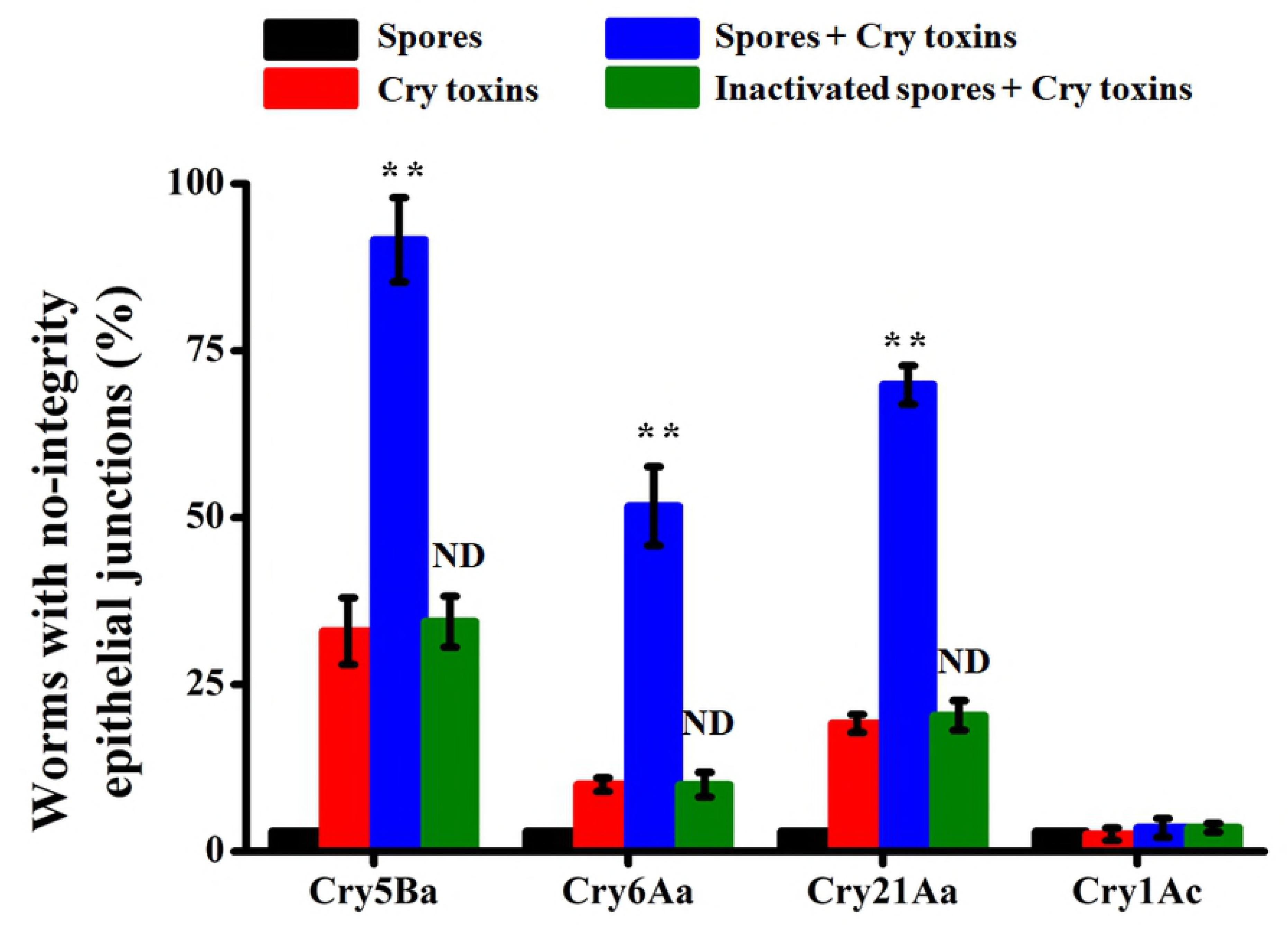
The presence of live-spores together with nematicidal Cry toxins are required for the disruption of the nematode epithelial junctions. The data are expressed as the percentage of worms with disrupted epithelial junctions after each treatment. The mean and standard error of three independent experiments are shown. Asterisks indicate significant differences (Student’s *t*-test, **, *p*<0.01) and ND indicates no significant difference between these treatments compared with that of corresponding pure Cry toxin treatment.

In another control we fed FT63(DLG::GFP) worms with a non-nematicidal Cry1Ac toxin, or we used inactivated spores from BMB171 strain together with Cry1Ac toxin. The epithelial junctions of most of the worms showed no disruption phenotype of their epithelial junctions (Fig. 6). Taking together, these results suggest that the developing of spores together with the action of a nematicidal Cry toxin are required for Bt to disrupt the nematode epithelial junctions. Therefore, it is possible that during the spore germination into vegetative cells *in vivo*, the nematicidal Bt strain would express additional virulence factors, besides the Cry toxins, resulting in the disruption of the epithelial junctions of the midgut tissue in the nematodes.

### The quorum sensing regulator PlcR plays important role in Bt disrupting nematode epithelial junctions

The quorum sensing regulon PlcR is responsible for the expression of additional Bt virulence factors to infect the hosts [19], including degradative enzymes, and enterotoxins [20,36,37]. On the other hand, the NprR regulator controls the necrotrophic properties of the bacteria to allow them to survive in the host cadaver [18,38] by regulating the expression of a set of genes encoding degradative enzymes [21]. In view of the importance of these regulons for the pathogenicity of Bt, we investigated whether PlcR or NprR play a role in the disruption of the epithelial junctions of the nematode.

The nematicidal Bt BMB0215 was used as the parental strain since this strain has the highest ability to disrupt the epithelial junctions of worms (Fig. 4), and compared it with the NprR-deficient (Δ*nprR*) and the PlcR-deficient (Δ*plcR*) strains, respectively. When these strains were used to fed FT63(DLG::GFP) worms, the percentage of destroyed epithelial junction caused by Δ*plcR* mutant strain was significant reduced (*t*-test, *p* < 0.05) than that of the parental strain (Fig. 7A). In contrast, a similar effect in the epithelial junctions was observed with the Δ*nprR* mutant strain compared with the parental strain (Fig. 7A). These results indicate that NprR does not participate in the ability of Bt to disrupt the epithelial junctions of the nematodes.

**Fig. 7.**
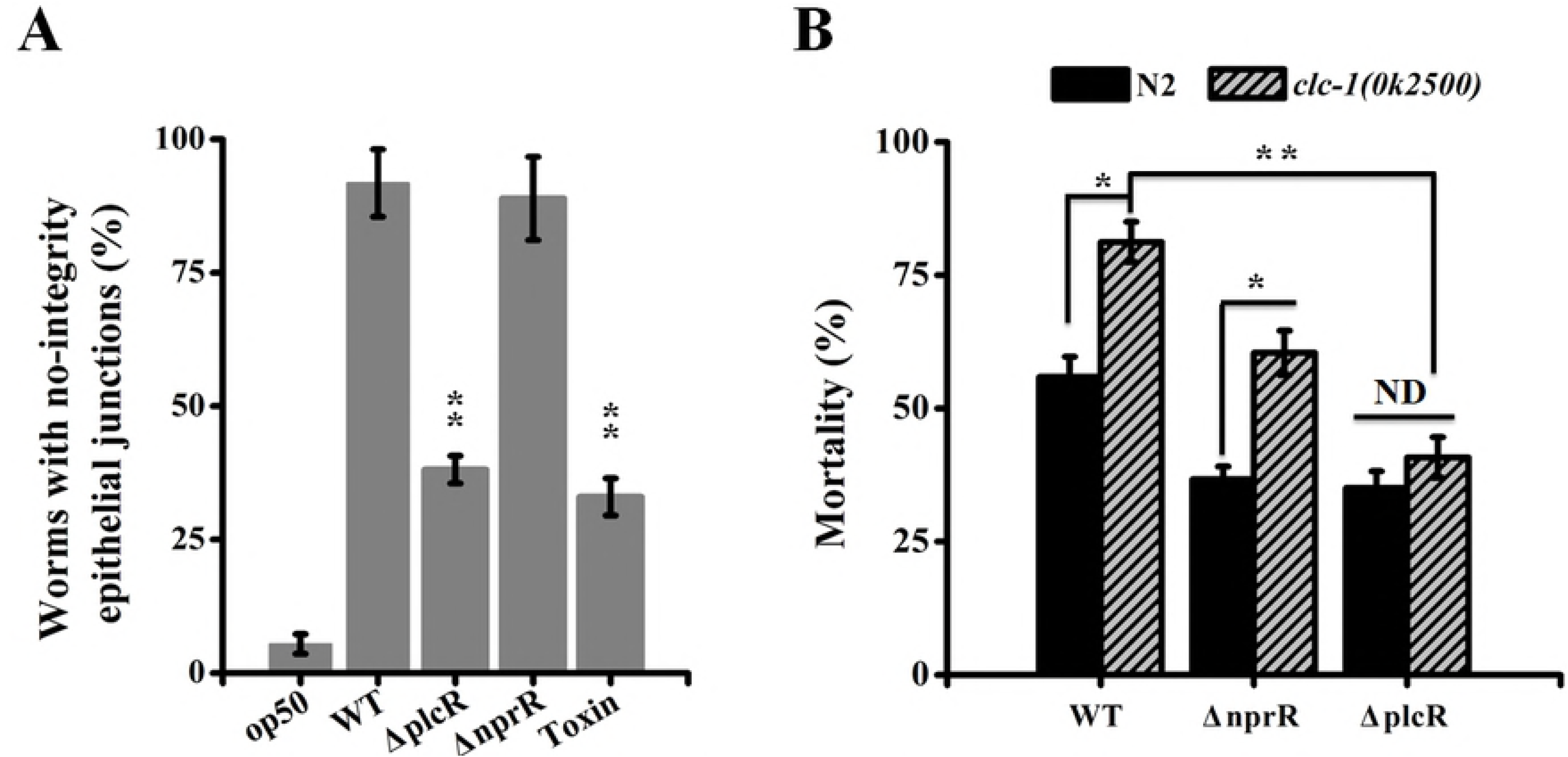
The PlcR regulon plays an important role in the disruption of the epithelial junctions of the worms induced by Bt bacteria. (A) Analysis of the disruption of epithelial junctions caused by *plcR* or *nprR* deficient Bt strains. The data are expressed as the percentage of worms that showed disrupted epithelial junctions after 24 h treatment for each strain. The mean and standard errors of three independent experiments are shown. (B) Toxicity assays of *plcR* or *nprR* deficient Bt strains against the wild type worms N2 or against the epithelial junction mutant worms *clc-1(ok2500)*. The mean and standard error of three independent experiments are shown. Asterisks indicate significance as determined by a Student’s *t*-test (*, *p*<0.05; **, *p*<0.01) and ND indicates no significant differences.

We also compared the nematicidal activity of the parental Bt strain with that of the Δ*nprR*, and Δ*plcR* mutant stains against the wild type N2 worms and the mutant *clc-1(ok2500)* worms that are defective in their epithelial junctions. Both the wild type N2 and mutant *clc-1(ok2500)* worms were more resistance to the infection with Δ*plcR* and Δ*nprR* Bt strains compared with that of the BMB0215 parental strain (*t-*test, *p*<0.05) (Fig. 7B). These results supported that both regulons PlcR and NprR controlled the expression of virulence factors that contribute to the Bt pathogenicity [18]. Even more, the parental BMB0215 strain and Δ*nprR* mutant Bt strain showed significant higher toxicity against the mutant worms *clc-1(ok2500)* than to the wild type worms N2 (*t-*test, *p*<0.05). However, the mortality was no significantly different for the Δ*plcR* mutant strain infecting the N2 and the epithelial junction defective mutant worms *clc-1(ok2500)* (*t-*test, *p*>0.05). Taken together, our data demonstrate that the quorum sensing regulator PlcR plays a central role disrupting the epithelial junctions in the worms.

### Identification of two PlcR-regulated virulence factors, ColB and Bmp1, that contribute to the epithelial junctions disruption induced by Bt strains

The role of PlcR in disrupting epithelial junctions is likely due to the activation of expression of specific virulence factors that targeted the intestinal epithelial junction of the worms. We previously described two metalloproteinase, named Bmp1 [23] and ColB [22], which are regulated by PlcR and are involved in the destruction of the intestine structure of *C. elegans*. Therefore, we investigated whether these virulence factors were involved in the disruption of the intestinal epithelial junction of worms induced after Bt infection. We fed FT63(DLG::GFP) worms with the parental Bt BMB0215, or the ColB-deficient (Δ*colB*), or the Bmp1-deficient (Δ*bmp1*), or the double ColB, Bmp1-deficient mutant (Δ*colb*Δ*bmp1*) strains. The results showed that the FT63(DLG::GFP) worms treated with either of the Δ*colB* or Δ*bmp1* single mutant strains had no significant differences in the disruption of epithelial junctions induced by the wild type BMB0215 strain (Fig. 8A). In contrast, when worms were fed with the double Δ*colb*Δ*bmp1*mutant the FT63(DLG::GFP) worms showed that the percentage of destroyed epithelial junction was significant lower (*t*-test, *p*<0.05) than that of the parental BMB0215 strain (Fig. 8A).

**Fig. 8.**
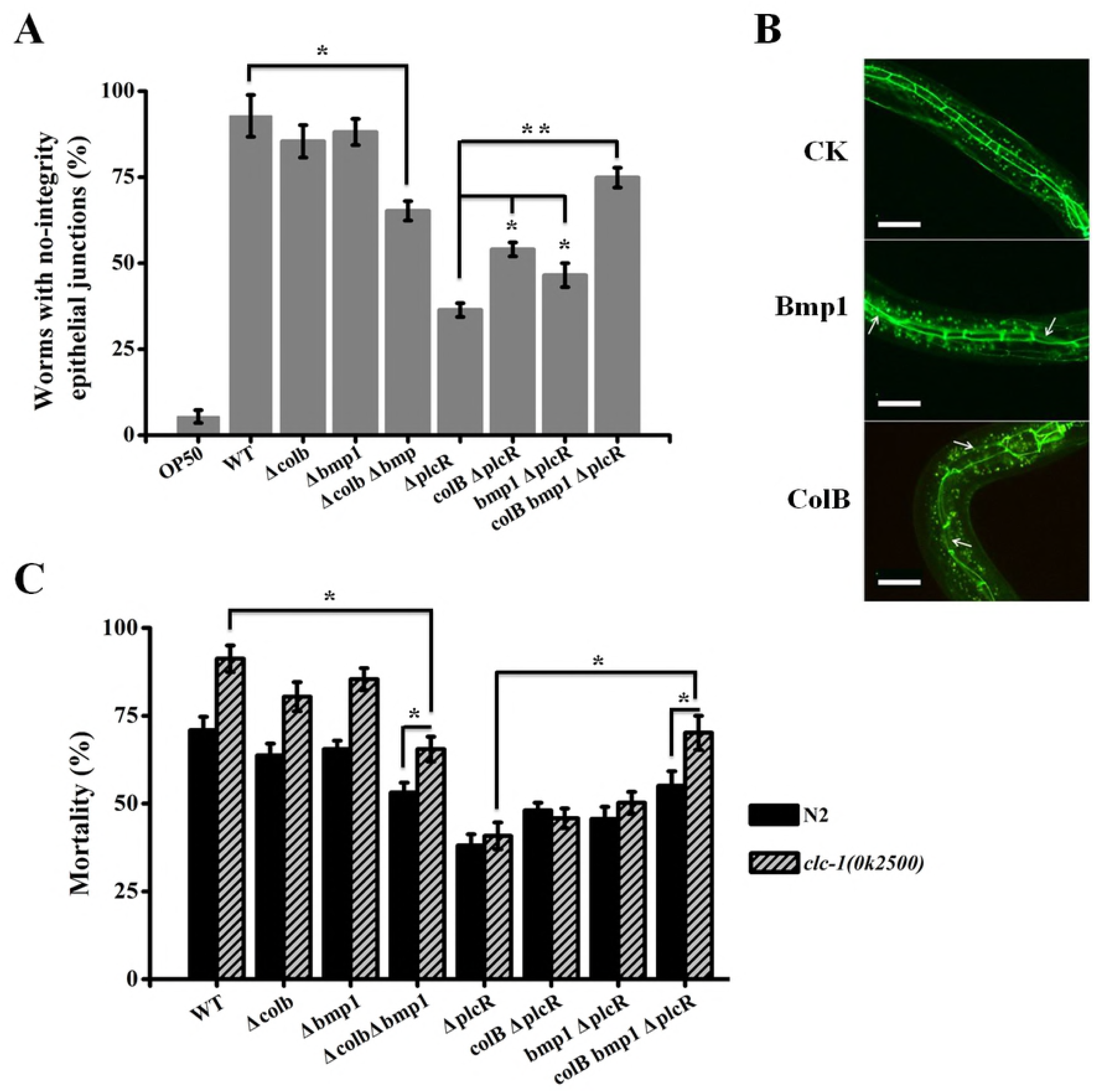
The PlcR regulated virulence factors ColB and Bmp1 contribute in the disruption of the epithelial junction of the worms induced by Bt. (A) Analysis of the disruption of epithelial junctions caused by different Bt mutant strains. The data are expressed as the percentage of worms that with no-integrity epithelial junctions after treatment. The mean and standard error of three independent experiments are shown. (B) Epithelial junction disruption observations of worms fed with *E. coli* strains over-expressing either ColB or Bmp1 metalloproteinases. Representative images of are shown. The scan bars represent 20 μm and the arrows indicate the disrupted of epithelial junctions. (C) The toxicity assay of mutant Bt strains against the wild type worms N2 or the epithelial junction mutant worms *clc-1(ok2500)*. The mean and standard error of three independent experiments are shown. Asterisks indicate significance as determined by a Student’s *t*-test (*, *p*<0.05; **, *p*<0.01) and ND indicates no significant difference.

To further analyze the role of ColB or Bmp1 metalloproteinases, FT63(DLG::GFP) worms were fed with *E. coli* strains over-expressing either ColB or Bmp1 metalloproteinases showing a clear disruption of the epithelial junctions of FT63(DLG::GFP) worms (Fig. 8B), indicating that high doses of Bmp1 and ColB have the potential to disrupt nematode epithelial junctions. Therefore, we compared the activity of disruption of junctions of the parental strain BMB0215 strain by measuring the disruption of the epithelial junctions of the Δ*plcR* strain harboring the plasmids that over express ColB or Bmp1 metalloproteinases under the control of a PlcR independent promoter (kanamycin promoter *P*_*apha3′*_ [39]). When expression of either *P*_*apha3′*_*colB* or *P*_*apha3′*_*bmp1* was induced in the Δ*plcR* mutant, the disruption of epithelial junctions was no significantly different to the Δ*plcR* strain. However, when both *P*_*apha3′*_*colB* and *P*_*apha3′*_*bmp1* were expressed together in the Δ*plcR* strain the activity of disruption of epithelial junctions was partially restored (Fig. 8A).

The toxicity of these strains against the mutant worms *clc-1(ok2500)* affected in their epithelial junctions confirmed the above conclusion (Fig. 8C). Both the wild type worms N2 and mutant worms *clc-1(ok2500)* were more resistance to the Bt Δ*colb*Δ*bmp1* double mutant strain compared with that of the parental strain BMB0215 (*t-*test, *p*<0.05) (Fig. 8C), supporting that ColB and Bmp1 contribute to the pathogenicity of Bt [22,23]. The Δ*colb*Δ*bmp1* double mutant strain showed significant higher toxicity against the mutant worms *clc-1(ok2500)* than that of the wild type N2 worms (*t-*test, *p*<0.05), although the mortality was no significantly different when either single mutant strain was used to infect the N2 and mutant *clc-1(ok2500)* worms (*t-*test, *p*>0.05) (Fig. 8C). Also, the over expression of *P*_*apha3′*_*colB* or *P*_*apha3′*_*bmp1* in the Bt Δ*plcR* mutant showed no significant difference in nematicidal activity when compared with the Δ*plcR* strain. But the nematicidal activity was partially restored when both metalloproteinases *P*_*apha3′*_*colB* and *P*_*apha3′*_*bmp1* were over expressed together in the Δ*plcR* strain (Fig. 8C). Thus, our results support that ColB and Bmp1 contribute to the pathogenicity of Bt via disrupting the epithelial junction of worms.

## DISCUSSION

In this study, we selected to work with different Bt strains in the nematode *C. elegans* as a model system to investigate the pathogenesis of Bt in natural conditions including virulence, necrotrophic, and sporulation stages. Our data showed that Bt bacteria disrupt the epithelial junctions of the nematode intestinal cells during its host invasion (Fig. 2 and Fig. 3). Moreover, we showed that targeting epithelial junctions is a widely adopted strategy since all the nematicidal Bt strains analyzed disrupted the epithelial junctions of intestinal cells in *C. elegans* (Fig. 3). Cell-cell junctions play a critical role in cell polarity and organogenesis, and constituted a physical barrier against pathogens invasion [35]. This functional barrier is composed of three major components, tight junctions (TJ), adherence junctions (AJ) and GAP junctions [35]. Several pathogens disrupt epithelia junctions to adhere and invade the deeper tissues for their proliferation [5]. For example, the *P. gingivalis*, a human adult pathogen, produces cysteine proteases to degrade the E-cadherin protein that is located at the AJ, facilitating the invasion of the underlying tissues through a paracellular pathway [7]. In the case of *C. elegans* it was shown that knockdown of either *dlg-1* or *ajm-1* genes, which encode intestinal epithelial junction components related protein DLG-1 or AJM-1 respectively, reduces worm lifespan upon *Enterococcus faecalis* challenge [34]. To our knowledge, this is the first study explaining the participation of epithelial junctions in *C. elegans*-Bt interactions.

In addition, we showed that the quorum sensing regulator PlcR of Bt, that is the main virulence regulator [19] required for the early steps of the nematode infection process [20,36,37], while the NprR regulator that controls the necrotrophic properties of the bacteria by regulating a set of genes encoding degradative enzymes [18,38] did not participate in the disruption of the epithelial junction of *C. elegans* (Fig. 7 and Fig. 8). Therefore, our data suggest that destroying host epithelial junctions is an early event in Bt pathogenesis that happens before the nematode death, rather than in the saprophytic stage. These data gave novel insight into the pathogenesis of Bt, since the epithelial junctions are conserved structures found in nematodes, insects, or mammals [35], suggesting that epithelial junction disruption could be a common mechanism among different Bt-host interactions.

However, although we show that PlcR plays an important role in the disruption of the nematode epithelial junctions (Fig. 7), it is undeniable that when *plcR* gene was deleted, there were about 35% worms that still showed disrupted epithelial junctions (Fig. 7A). There may be two possible explanations: the first is that other virulence factors not controlled by the PlcR regulon could also target the epithelial junction. The second, is that the nematicidal Cry toxin may also target the epithelial junctions of *C. elegans*, since treatment with a purified Cry toxin without Bt bacteria also lead to some damage of the epithelial junctions from 10.0 ± 1.0 % for Cry6Aa to 33.0 ± 5.0% for Cry5Ba (Fig. 6). It was previously shown that AJ of mammalians cells participate in the susceptibility to another pore-forming toxin, the α-toxin produced by *Staphylococcus aureus* bacteria [40]. Thus, it could be possible that the Cry toxin may be a key factor to initiate the process of disruption of the nematode epithelial junctions. Purified Bt spores alone failed to germinate [22] and did not disrupt the epithelial junctions, while live spores together with the nematicidal Cry toxin act synergist to disrupt the nematode epithelial junctions (Fig. 5). Moreover, the specificity of Cry toxins also plays a crucial role in this process, since the effect on the epithelial junctions was not observed when the nematodes were treated with the lepidopteran specific Cry1Ac toxin and Bt spores (Fig. 6). The specific binding of Cry toxins to their corresponding receptors may determine the pore formation activity of the toxin and whether Bt can effectively initiate the disruption of the epithelial junctions in the susceptible host. This strategy may be the result of long-term co-evolution between Bt and its host.

Regarding the virulence factors directly involved in the disruption of epithelial junctions, our data show that at least *colB* and *bmp1* genes that are activated by the PlcR regulon are involved in this process. However, other virulence factors participate since the deletion of *colB* or *bmp1* genes alone in its parental strain did not reduce the phenotype of epithelial junction disruption (Fig. 8A). Only the double knockout mutant of *colb* and *bmp1* genes resulted in a reduced disruption of epithelial junctions, that was lower than the PlcR mutant strain (Fig. 8B). Since the PlcR regulon controls the expression of various virulence factors, our results indicate that there are other virulence factors regulated by PlcR that are involved in this process.

A model explaining the Bt induced disruption of the intestinal epithelial junction in *C. elegans* is shown in Fig. 9. The spores and toxins of nematicidal Bt are ingested by the worms, the toxins bind to specific receptors on the midgut epithelial cells, forming pores in the intestinal cell membranes, creating favorable conditions for the spore germination and bacterial growth [8], vegetative cells express virulence factors mainly controlled by the PlcR regulon inducing the disruption of the worm’s epithelial junctions at the early stage, and leading to the decomposition of the intestinal cells. The tightness of the intestinal cells collapses, making a pathway for vegetative cells to pass through the intestine and invade the worms. Finally, the Bt cells express additional virulence factors controlled by NprR regulon that allow the reproduction of Bt cells on the nutrient source of the cadaver leading to the release of spores into the environment [17,41].

**Fig. 9.**
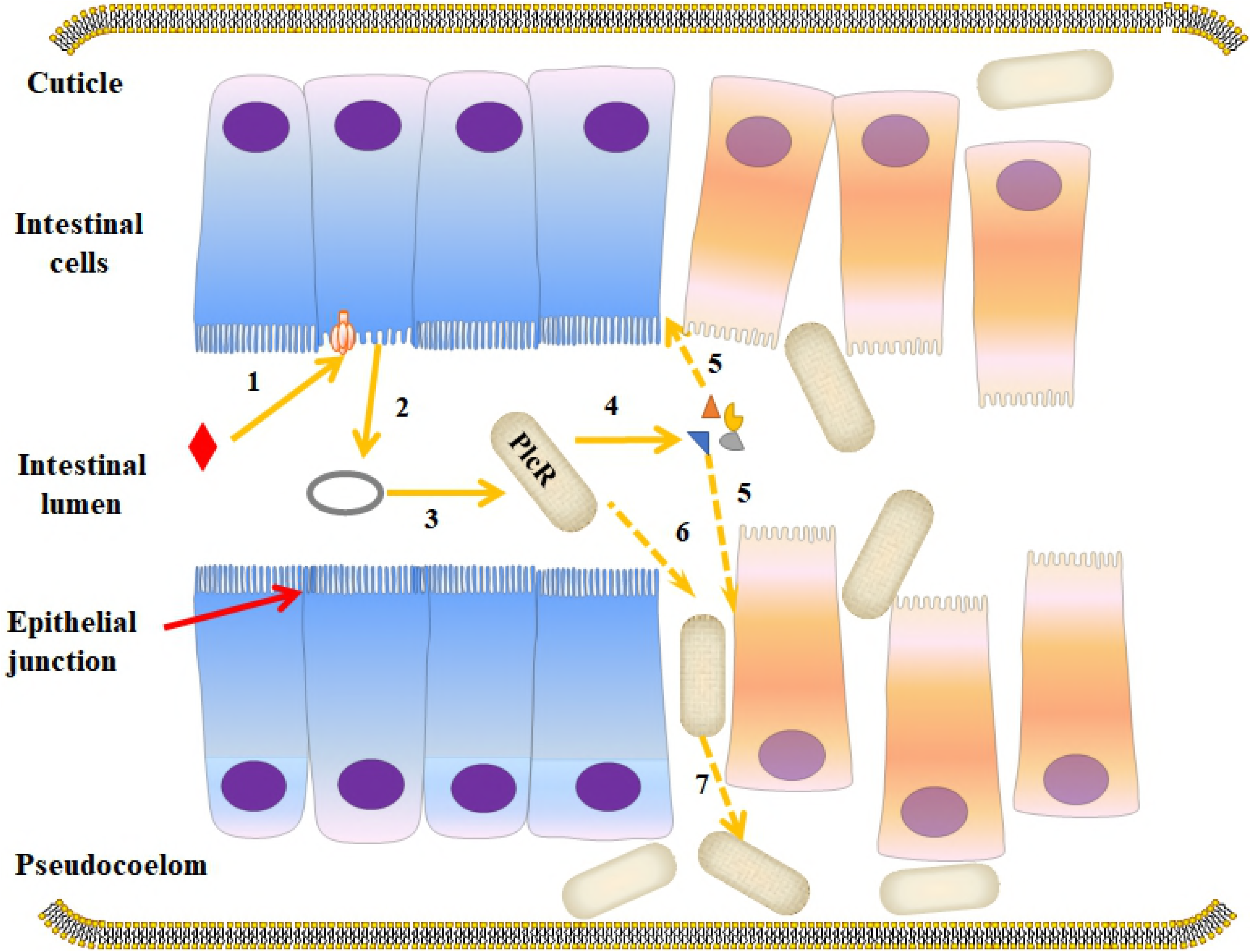
Model of Bt infection cycle by disrupting the epithelial junction of nematodes. After the nematicidal Bt spore/Crystals were fed to the worms, the Bt spores are resistant to the pharyngeal grinder and both spores and Cry toxins reach the midgut lumen. The Cry toxins form pores in the membrane of the midgut epithelial cells (1), leading to the germination of spores in the intestinal lumen (2). The spores then could develop into vegetative cells (2), which express different virulence factors of the PlcR regulon. The various irregular shapes represent the virulence factors involved in Bt to disrupting the host epithelial junctions such as Bmp1 and ColB, and other PlcR-independent factors (4), that participate in the disruption of epithelial junctions of worms (5). Then the vegetative cells could break through the epithelial cells (6), multiply in the host and collapse the intestinal lumen of the worms (7). The solid arrows indicate steps that have been described before. The dashed arrows represent the steps described in this study.

One of the most studied nematicidal Bt toxin is Cry5B, which specifically binds to the glycolipid receptor located in the apical membrane of the intestinal epithelial cells, leading to insertion of the toxin into the plasma membrane and pore formation [42]. Worms respond to Cry5B intoxication by inducing a conserved innate immunity response including the activation of the mitogen-activated protein kinase (MAPK) pathways p38/MAPK and c-Jun/MAPK [43], the endoplasmic reticulum unfolded protein response (UPR) pathway [43,44], the hypoxia response pathway and the DAF-2 insulin/insulin-like growth factor-1 signaling pathway [45,46]. In this study, we describe that PlcR regulated virulence factors help Bt to infect *C. elegans* by disrupting its epithelial junctions (Fig. 7 and Fig. 8), suggesting that epithelial junctions serve as a physical barrier to protect *C. elegans* against the infection of pathogens such as Bt. *C. elegans* is a soil-dwelling and free-living animal [47], which encounters a myriad of chemical or biological hazards in the environment. The cuticle naturally serves as the first physical barrier to protect the nematodes from these detrimental agents [48,49]. As *C. elegans* feeds on bacterial cells, the pharyngeal grinder should serve as a second barrier to break up bacteria and preventing them to contact their intestine [50]. However, Bt spores are resistant to the pharyngeal grinder [51]. Thus, the intestine might be the last physical barrier that prevents the spore germination into vegetative bacteria in the invaded worms. Therefore, we propose that the epithelial junction serves as the crucial connection of intestinal epithelial cells to prevent the invasion of Bt pathogen in *C. elegans*. Our work highlights the importance of host epithelial junctions as a physical barrier in the pathogen-host interactions and describe novel mechanisms used by Bt to successful invasion and worm infection.

## MATERIALS AND METHODS

### Bacterial strains, plasmids and culture conditions

The bacterial strains and plasmids used in this study are listed in Table S1. All *Escherichia coli* and Bt strains were grown on Luria-Bertani (LB) medium supplemented with the appropriate antibiotics at 37 °C or 28 °C, for *E. coli* or Bt, respectively.

### *Caenorhabditis elegans* strains and culture conditions

The wild-type *C. elegans* strain N2, the epithelial junction GFP labeled transgenic strain FT63 (DLG::GFP), and the epithelial junction mutant strain *clc-1(ok2500)* were provided by the *Caenorhabditis* Genetics Center (CGC, http://www.cbs.umn.edu/CGC/). Worms were cultured at 20 °C on NGM plates (0.3% NaCl, 0.25% tryptone and 1.5% agar) with *E. coli* strain OP50 as food source [47]. The synchronized L4 stage worms were prepared according to a previous described method [52].

### Preparation of the Cry proteins

For Cry protein production the Bt strains described in table S1 were cultivated in liquid ICPM medium (0.6% tryptone, 0.5% glucose, 0.1% CaCO_3_, 0.05% MgSO_4_ and 0.05% K_2_HPO_4_, PH 7.0) at 28 °C, 220 rpm for three days until complete sporulation. The crystal inclusions were purified as previously described [53].

### Preparation and inactivation of the spores

The spores of an acrystalliferous mutant of *B. thuringiensis* strain BMB171 were obtained by culturing cells in CCY sporulation medium [54] at 28 °C for 4 days. Spores were harvested by centrifugation of the cultures at 10,000 *xg* for 15 min at 4 °C and washed 3 times with sterile distilled water (50 ml). The spore pellets were suspended in 10 ml sterile distilled water and store in −20 °C. The purified spores were activated at 70 °C for 15 min and induced for germination in LB medium at 37 °C for 1 h. Germinated spores were then inactivated by heat treatment at 80 °C for 15 min. The surviving spores were then induced to germinate in LB medium again. The germination and inactivation procedures were repeated twice to make sure that above 99% spores were inactivated.

### RNA extraction and qRT-PCR

Total RNA was extracted from *C. elegans* using TRIzol reagent (Invitrogen, Carlsbad, California, USA). The cDNA was reverse transcribed with random primers using Superscript II reverse transcriptase (Invitrogen, Carlsbad, CA, USA) according to the manufacturer’s protocol. The expression analyses of certain genes in *C.elegans* were performed using qPCR with the primers listed in Table S2. The qPCR was conducted with Life Technologies ViiA™ 7 Real-Time PCR system (Life Technologies, California, USA) using the Power SYBR Green PCR Master Mix (Life Technologies, CA, USA) according to the manufacturers’ instructions. The experiments were conducted in triplicate. Primer efficiency correction was used in 2^-ΔΔ^Ct relative quantitation analysis using *tba-1* gene for reference and normalization.

### Cloning of Cry21Aa gene

The primers F-Cry21Aa and R-Cry21Aa (Listed in Table S2) were designed based on the genome sequence of C15 strain. These primers were used to amplify *cry21Aa* gene including its promoter and terminator. The 5.5-kb PCR fragment was purified and cloned into the *Bam*HI*-Hind*III site of the *E. coli*-Bt shuttle vector pHT304 to generate recombinant plasmid pHT304-cry21Aa. The recombinant plasmid was confirmed by DNA sequencing (AuGCT, Beijing, China) and transferred into Bt strain BMB171 to generate recombinant strain BMB171/Cry21Aa for expression Cry21Aa protein.

### Construction of Bt knockout mutant strains

DNA fragments of Bt BMB0225 strain corresponding to the upstream and downstream regions of the target genes (described in results section) were generated by PCR using the primers listed in Table S2. The two amplified fragments were digested with *Bam*HI-XbaI and *Bam*HI-*Kpn*I, respectively, and cloned into the temperature-sensitive plasmid pHT304-Ts. Then, the spectinomycin resistance gene from plasmid pBMB2062 was digested with *Bam*HI and inserted between these two gene fragments. The resulting plasmid was transformed into BMB0225 strain and cultivated for 8 h in LB medium supplemented with 25 μg/ml erythromycin. The transformant cells were cultivated at 42 °C for 4 days for selection of bacterial cells that have integrated the plasmid by homologous recombination. Erythromycin-resistant (25 μg/ml) but spectinomycin sensitive (20 μg/ml) colonies were selected. The mutant strains (Table S1) were confirmed by PCR and DNA sequencing (AuGCT, Beijing, China).

### Bag-of-bacteria (Bob) phenotype assay

The “Bob” formation bioassay was performed in liquid S-media on 96-well plates as previous described [30]. In this assay we used 100 µl mixture per well consisted of 80 µl S-media, 10 µl *E. coli* OP50 strain at OD_600_ of 0.3, 10 µl spores or spore/crystal mixtures (a 10^6^ dose of spores per well) and 30∼50 synchronized L4 stage worms in three independent replicates. The 96-well plates were incubated at 20 °C and worms monitored from 24 to 96 h for the formation of “Bob” phenotype under differential interference contrast (DIC) microscopy observations using 400 X magnification (Olympus BX51, Olympus, Tokyo, Japan).

### Analysis of *C. elegans* epithelial junctions

The epithelial junctions’ observation assays were performed in liquid S-media on 48-well plates. For various Bt strains we used 300 µl mixture per well consisted of 270 µl S-media, 20 µl *E. coli* OP50 strain at OD_600_ of 0.3, 10 µl spores or spore/crystal mixtures (10^6^ dose of spores per well), and approximately 200 synchronized L4 stage FT63(DLG::GFP) worms. For various Cry proteins we used 300 µl mixture per well consisted of 270 µl S-media, 20 µl *E. coli* OP50 strain at OD_600_ of 0.3, 10 µl spores (10^6^ dose of spores per well) or spore/Cry toxin mixtures (10^6^ dose of spores plus 10 ng/µl Cry toxins), and approximately 200 synchronized L4 stage FT63(DLG::GFP) worms. The 48-well plates were incubated at 20 °C and worms monitored after 24 h of treatment Observations were performed under fluorescent microscope (Olympus BX51, Olympus, Tokyo, Japan) at 100X magnification after 18 h of treatment. The percentages of worms with disrupted epithelial junctions after each treatment were calculated based on the observation of at least 100-120 individual worms.

The transformants *E. coli* BL21(DE3) cells expressing the proteinases ColB [22] and Bmp1 [23], were grown at 37 °C to the mid-log phase (OD_600_ 0.4 to 0.6) and then 0.1 mM IPTG (final concentration) was added to induce the expression of these virulence factors after incubation at 28 °C for additional 6 h. Then bacteria were spread onto NGM plates (0.3% NaCl, 0.5% tryptone, 0.1% yeast extract and 1.5% agar) containing 0.1 mM IPTG and cultured overnight at 28 °C. About 200 synchronized L4 stage FT63(DLG::GFP) nematodes were placed onto each plate incubated at 20 °C, and monitored for the disruption of epithelial junctions after 8 h of treatment. *E. coli* BL21(DE3) cells harboring empty vector were used as negative control under the same conditions. The percentage of worms that showed disrupted epithelial junctions after each treatment was calculated based on the observation of at least 100-120 individual worms. All tests were conducted in three independent replicates.

### Nematode mortality assays

Bt strains were grown at 28 °C in LB medium overnight and spread onto NGM plates at 28 °C for 4∼6 days until the spore/crystal mixture covered the plates. Then 20∼30 L4 stage synchronized worms were transferred onto these NGM plates spread with each Bt lawn as the only food source for the worms that feed on spore and crystal mixtures freely. The plates were incubated at 20 °C and mortality was recorded after 3 days. We used the food source strain *E. coli* OP50 and nematicidal Bt strain BMB0215 as negative and positive controls, respectively. All tests were conducted with three independent replicates and monitored for survival at indicated time.

### Microscopic observations

Worms were photographed on 2% agarose pads using the fluorescence microscope and the phase contrast microscope (Olympus BX51, Olympus, Tokyo, Japan).

### Data analysis

The data were analyzed using Statistical Package for the Social Sciences, (SPSS version 13.0 Chicago, IL, USA). Statistic comparisons used a Student’s *t* test and significant differences were determined as a threshold of *p*<0.05 (*), *p*<0.01 (**), or *p*<0.001 (***).

## ACKNOWLEDGMENTS

We thank professor Didier Lereclus (INRA, Guyancourt, France) for kind donation of plasmids pHT315-*papha3′gfp* and pHT315. We also thank the Caenorhabditis Genetics Center for worm strains.

## Supporting Information Legends

Table S1. Bacterial strains and plasmids used in this study.

Table S2. Primers used in this study.

